# Putative TetR family transcriptional regulator Rv1019 of *Mycobacterium tuberculosis* is an auto-repressor and a negative regulator of *mfd-mazG* operon

**DOI:** 10.1101/2020.02.10.941963

**Authors:** Akhil Raj Pushparajan, Ranjit Ramachandran, Jijimole Gopi Reji, Ramakrishnan Ajay Kumar

## Abstract

Regulation of gene expression at the level of transcription is one of the ways living organisms differentially make proteins in their cells. Intracellular pathogen *Mycobacterium tuberculosis* persists in the host for years in a latent state in structures called granuloma. Rv1019, a member of an uncharacterized TetR family of transcriptional regulators of *Mtb* H37Rv, was found to be differentially expressed during dormancy and reactivation *in vitro*. By GFP reporter assays, electrophoretic mobility shift assays and chromatin immuno-precipitation assays we showed that this protein binds to its own promoter and acts as a negative regulator of its own expression. It forms dimers *in vitro* which is essential for the DNA binding activity, which is abrogated in the presence of tetracycline. We show that *Rv1019* and downstream genes *Rv1020* (*mfd*), *Rv1021* (*mazG*) are co-transcribed. Constitutive expression of *Rv1019* in *M. smegmatis* downregulated *MSMEG_5423* (*mfd*) and *MSMEG_5422* (*mazG)* suggesting that Rv1019 negatively regulates these downstream genes. *M. smegmatis* expressing *Rv1019* was found to be sensitive to UV and H_2_O_2_ compared to the control suggesting its role in regulating DNA damage response in mycobacterium.

**Importance:** One of the reasons for our inability to eradicate tuberculosis, in spite of the availability of effective drugs and BCG vaccine, is the ingenuity of *Mycobacterium tuberculosis* to survive, persist and multiply inside the lungs where it employs clever strategies to counter the host defence systems. The number of anti-TB drugs in the current therapeutic regimen is limited, and the long duration of treatment increases the chances for emergence of drug resistant bacteria. Rv1019, a transcriptional regulator differentially expressed in *in vitro* dormancy-reactivation studies, was found to be an auto-repressor and negatively regulating *mfd-mazG* operon involved in DNA repair. As Rv1019 is a regulator of critical pathways in bacterial persistence, it could be a prospective target for prophylactic intervention.

## Introduction

Tuberculosis (TB), caused by *Mycobacterium tuberculosis* (*Mtb*), is one of the oldest diseases and is the leading cause of death of humans by an infectious agent. Ten million people have been estimated to be infected with *Mtb* causing approximately 1.2 million death annually (WHO, 2019). *Mtb* has the ability to adapt and persist in the human body in a dormant state in structures called garnuloma for long periods without manifesting any clinical symptoms (1). The ability of *Mtb* to remain dormant is the major reason for the ineffectiveness of TB eradicating programs, longer duration of treatment and emergence of drug resistance (2).

Availability of complete sequence of *Mtb* genome has led to a better understanding of the pathogen for devising therapeutic strategies (3, 4). *Mtb* genome codes for approximately 4000 proteins of which 214 are transcriptional regulators which regulate expression of genes at the level of transcription. Many of them form homodimers and bind the regulatory regions to activate or repress the expression of downstream genes such as those of the two-component regulators PhoP, DevR etc (5, 6). TetR family of transcriptional regulators belongs to one-component prokaryotic signal transduction systems containing a helix-turn-helix DNA binding domain at the N-terminal and a C-terminal ligand binding domain. They are often repressors which usually bind around the RNA polymerase binding regions and regulate the expression of efflux pumps in bacteria (7). Binding of ligands at the C-terminal end causes a conformational change in the N-terminal region and abolish the DNA binding activity, thus relieving the repression (8). They bind incomplete palindromic region upstream to their own coding region or a region upstream of the regulated region (9, 10). TetR family of transcriptional regulators controls a diverse function including multi-drug resistance (11, 12), morphogenesis and antibiotic production (13, 14), osmotic stresses (15), catabolic pathways (16, 17), modification and clearance of toxic compounds (14) and virulence (17, 18).

Rv1019 is a transcriptional regulator of TetR family of *Mtb* H37Rv that was found to be differentially expressed in *in vitro* dormancy-reactivation studies (19, 20). In the present study we made an effort to characterize the protein by analysing its regulatory mechanism in *Mtb*. Our results show that similar to other TetR family of transcriptional regulators, Rv1019 is an auto-repressor and it negatively regulates the expression of downstream genes which are involved in DNA damage response.

## Results

To study the role of Rv1019 in *Mtb*, we amplified its ORF from *Mtb* H37Rv genome with gene specific primers **(Fig. S1 A)**. Amplified product was then cloned into pET32-a expression system and expressed in *E. coli* BL21 (DE3) cells, and the recombinant protein was purified by Ni-NTA immobilized metal affinity chromatography (IMAC) **(Fig. S1 B)**. The identity of the protein was confirmed by western blot using histidine antibodies **(Fig. S1 C)**. Polyclonal antibodies against the recombinant protein were generated in New Zealand White Rabbits, and purified. Specificity of the antibodies was confirmed by western blot **(Fig. S1 D)**.

### Rv1019 is an auto-repressor and a negative regulator of its own expression

Rv1019 belongs to TetR family of transcriptional regulators which are generally auto-repressors. To study the regulatory activity of Rv1019 on its own promoter, we first generated a knockout strain of *M. smegmatis* mc^2^155 which lacks *MSMEG_5424*, the homologue of *Rv1019* **(Fig. S2 A)**. Further we created a series of GFP reporter constructs with pFPV27 as the backbone plasmid (Fig. 1A, **S2 B)**. All the constructs were then electroporated into the *M. smegmatis ΔMSMEG_5424* strain and the GFP expression was monitored. By GFP analysis we found that *Rv1019* promoter is active in *M. smegmatis* and the strain harbouring promoter with *Rv1019* ORF (Construct IV) showed significant reduction in GFP expression compared to bacteria carrying construct III, the plasmid containing only the promoter (Fig. 1B, **C)**. For further validation of auto-repression, *Rv1019* promoter was cloned upstream of GFP and into the same construct, we subcloned *Rv1019* ORF under the *hsp60* promoter in the opposite orientation (Construct V) (Fig. 1A). Interestingly *M. smegmatis* with these constructs showed a significant reduction in fluorescence when compared to that carrying the promoter-GFP alone controls (Fig. 1D). These results confirmed that Rv1019 binds to its own promoter and represses the expression of its own gene. This was further confirmed by analysing GFP expression by confocal microscopy **(Fig. S2 C)**. To study the *in vivo* binding, we performed a chromatin immunoprecipitation (ChIP) on the lysate of actively growing log phase *Mtb* with Rv1019 antibodies and checked the precipitated DNA for *Rv1019* promoter by PCR. We were able to amplify the *Rv1019* promoter but not the *hsp65* promoter, indicating that under normal aerobic laboratory growth conditions Rv1019 is recruited to its own promoter (Fig. 1E).

**Fig. 1.**
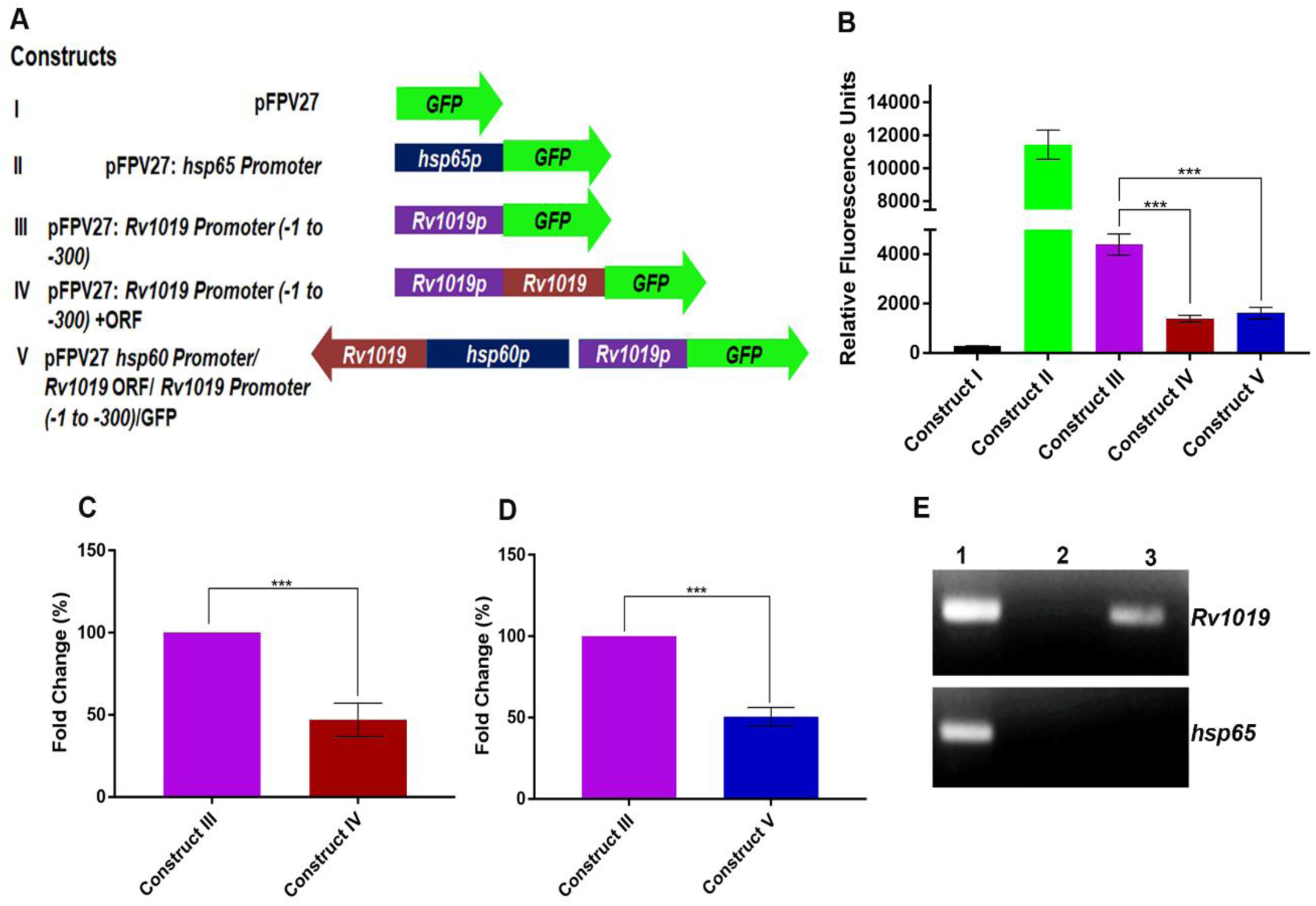
Rv1019 is an auto-repressor. **(A)** Auto-regulation of *Rv1019* expression was analysed in *M. smegmatis* with pFPV27 reporter system containing GFP. Construct I: promoterless pFPV27 vector; construct II: vector carrying *hsp65* promoter; Construct III: −1 to −300 region of *Rv1019* cloned upstream of GFP. Construct IV: Rv1019 ORF with its 300 bp upstream region was cloned as a transcriptional fusion upstream of GFP. Construct V: Rv1019 ORF under *hsp60* promoter in reverse orientation of GFP under −1 to −300 region of Rv1019 promoter. All the constructs were electroporated into *M. smegmatis ΔMSMEG_5424* (*MSMEG_5424* is the homologue of *Rv1019*) and the fluorescence was measure after 48 h and represented in terms of RFU. (**B)** GFP expression from construct III vs construct IV and construct III vs construct V were compared and values are represented as mean of three independent experiments (Student’s t-test). ***P ≤ 0.0001 and ***P ≤ 0.0002. (**C)** RFU of *M. smegma*tis carrying Construct IV against that of the bacterium carrying Construct III is represented as percentage, values are the mean ± standard deviation of three independent experiments (Student’s *t*-test).***P ≤ 0.0001. **(D)** RFU of recombinant *M. smegmatis* carrying Construct V against that of the bacterium carrying Construct III. The values are the mean ± standard deviation of three independent experiments (Student’s *t*-test). ***P ≤ 0.0008. **(E)** ChIP assay to demonstrate *in vivo* binding of Rv1019 to its own promoter in *Mtb* grown under standard laboratory conditions. Sheared *Mtb* DNA–protein complex was immunoprecipitated with antibodies against Rv1019, and the DNA in the immune complexes was analyzed by PCR using specific primers for *Rv101*9 and *hsp65* promoters. Lane 1: PCR amplicon of input DNA, Lane 2: PCR amplicon of IgG pulldown, Lane 3: PCR amplicon of Rv1019 pulldown.

### Rv1019 binds to an 18 bp incomplete inverted repeat in its own promoter

To study the binding of Rv1019 on its own promoter, we conducted electrophoretic mobility shift assay (EMSA) using ^32^P-labeled 300 bp upstream sequence (5 nM) with increasing concentrations of recombinant Rv1019 (2–10 μM). Formation of a distinct DNA-protein complex was observed intensity of which increased as a function of the concentration of the protein (Fig. 2A). Specificity of binding was proven by competitive and non-competitive EMSA. In competitive EMSA, labelled DNA in the complex was replaced by unlabelled DNA of the same sequence (10 and 100-fold excess), making the majority of the DNA in the complex nonradioactive and hence invisible during imaging. In contrast, binding was not affected even at 100-fold excess of nonspecific promoter (of an irrelevant *Mtb* gene *Rv0440c* that codes for GroEL2 (Fig. 2B).

**Fig. 2.**
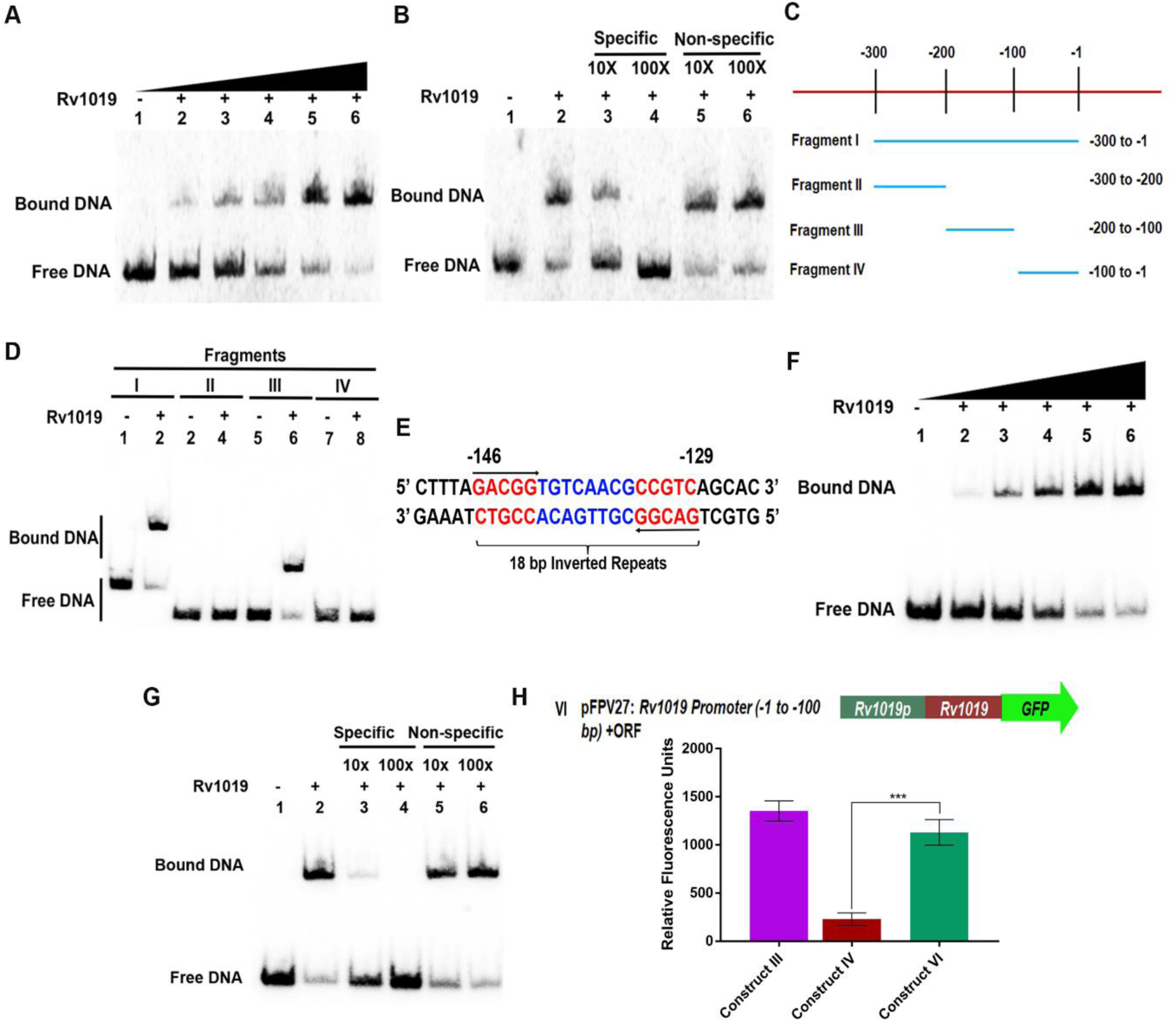
Rv1019 binds to an 18 bp inverted repeat upstream of its ORF. **(A)** EMSA shows *in vitro* binding of Rv1019 to −1 to −300 region upstream of *Rv1019* ORF. Lane 1: 1 nM ^32^P-labelled DNA without Rv1019, Lanes 2-6: increasing concentrations of Rv1019 (2-10 µM); **(B)** Competitive and non-competitive EMSA to validate the specificity of binding. Lane 1: ^32^P-labeled DNA without protein, Lane 2; ^32^P-labelled DNA incubated with 10 µM of Rv1019 (positive control), Lanes 3-4: competitive EMSA, Lanes 5-6 non-competitive EMSA; **(C)** Mapping of cognate-binding sequence of Rv1019 upstream of ORF; **(D)** EMSA with DNA fragments upstream of ORF. Lanes 1-2: −1 to −300 upstream, Lanes 3-4: −200 to −300 upstream, Lanes 5-6: −100 to −200 upstream, Lanes 7-8: −1 to −100 upstream. **(E)** 18 bp inverted repeat identified in −100 to −200 region by palindrome finder. Inverted repeats (red), gap (blue) and flanking region (black); **(F)** EMSA on 18 bp palindrome. Lane 1: ^32^P-labelled palindrome without Rv1019, Lanes 2-6: increasing concentrations of Rv1019; **(G)** Competitive and non-competitive EMSA to validate the binding on palindrome. Lane 1: ^32^P-labeled DNA without protein, Lane 2; ^32^P-labelled DNA incubated with 10 µm of Rv1019 (positive control), Lanes 3-4: competitive EMSA, Lanes 5-6 non-competitive EMSA; **(H)** GFP reporter construct to confirm the 18 bp binding region *in vivo.* Construct VI: −100 to −1 region with *Rv1019* ORF excluding the 18 bp cognate binding site was cloned upstream of GFP in pFPV vector. GFP expression was analyzed in *M. smegmatis ΔMSMEG_5424*; **(I)** GFP expression from construct IV vs Construct VI was compared. The values are the mean ± standard deviation of three independent experiments (Student’s *t*-test). ***P ≤ 0.0005.

To determine the exact binding site of Rv1019 on its promoter, PCR-amplified fragments labelled with [γ-^32^P] ATP were subjected to EMSA after incubating it with the recombinant protein (Fig. 2C). At 10 μM concentration of recombinant Rv1019, we observed formation of DNA-protein complexes as evidenced by a distinct shift in the migration pattern of DNA fragments I (–300 to 0) and III (–200 to −100) while the fragment spanning –300 to –200 (Fragment II) and −100 to 0 (Fragment IV) did not show any shift in the presence of the protein (Fig. 2D). These results suggested that the binding region is located between –200 to −100 from the start site of the *Rv1019*. Bioinformatic analysis showed the presence of an RNA polymerase binding site (−110 to −60 upstream of ORF) close to Rv1019 binding region as previously reported (21).

Many of the transcriptional regulators have been found to bind to inverted palindromic sequences upstream of the regulated genes. Further, *in silico* analysis of the −200 to −100 region upstream of the *Rv1019* revealed an 18 bp incomplete inverted repeat sequence (Fig. 2E). EMSA using this region as the template showed formation of DNA-protein complex (Fig. 2F). Binding was further validated with competitive and non-competitive EMSA (Fig. 2G). This was further confirmed *in vivo* employing a reporter construct which lacked the 18 bp promoter region (Construct VI) **(Fig. S3 A)**. Increased fluorescence of *M. smegmatis* carrying the construct in comparison with the strain carrying the construct which contained the cognate binding site confirmed that the protein binds to the 18 bp sequence leading to auto-repression (Fig. 2H and **Fig. S3 B)**.

### Rv1019 forms homodimers *in vitro*

Sequence analysis of Rv1019 at NCBI-CDD revealed the presence of an N-terminal helix-turn-helix motif which is conserved among DNA binding transcriptional regulators of *Mtb*. These proteins form dimers or higher order forms to bind to its target DNA (7). Therefore, we tested if Rv1019 could exist as a dimer *in vitro*.

Homology modelling of Rv1019 was performed in SWISS-MODEL server which revealed that the protein can exist as homodimers (Fig. 3A). To test dimer formation, the recombinant protein was electrophoresed on a polyacrylamide gel under non-denaturing conditions where we could see a clear band of 44 kDa at the expected size of a dimer compared to the size of the protein under denaturing condition confirming that protein can form dimers *in vitro* (Fig. 3B). Further, purified recombinant Rv1019 was covalently crosslinked by treating it with glutaraldehyde and then analysed by SDS/PAGE. In the absence of any cross-linking reagent, Rv1019 was seen as a monomer of 22 KDa, while in presence of 0.0001% and 0.0005% glutaraldehyde, the protein readily formed a homodimer of about 44 KDa (Fig. 3C). Bands seen above the dimer band probably represent multimers formed in the presence of glutaraldehyde. Western blot using antibodies against Rv1019 confirmed the identity of the monomer, dimers and higher order forms (Fig. 3D). These were further confirmed to be different forms of Rv1019 by excising the bands and subjecting them to MALDI-TOF/MS/MS analysis **(Fig. S4**).

**Fig. 3.**
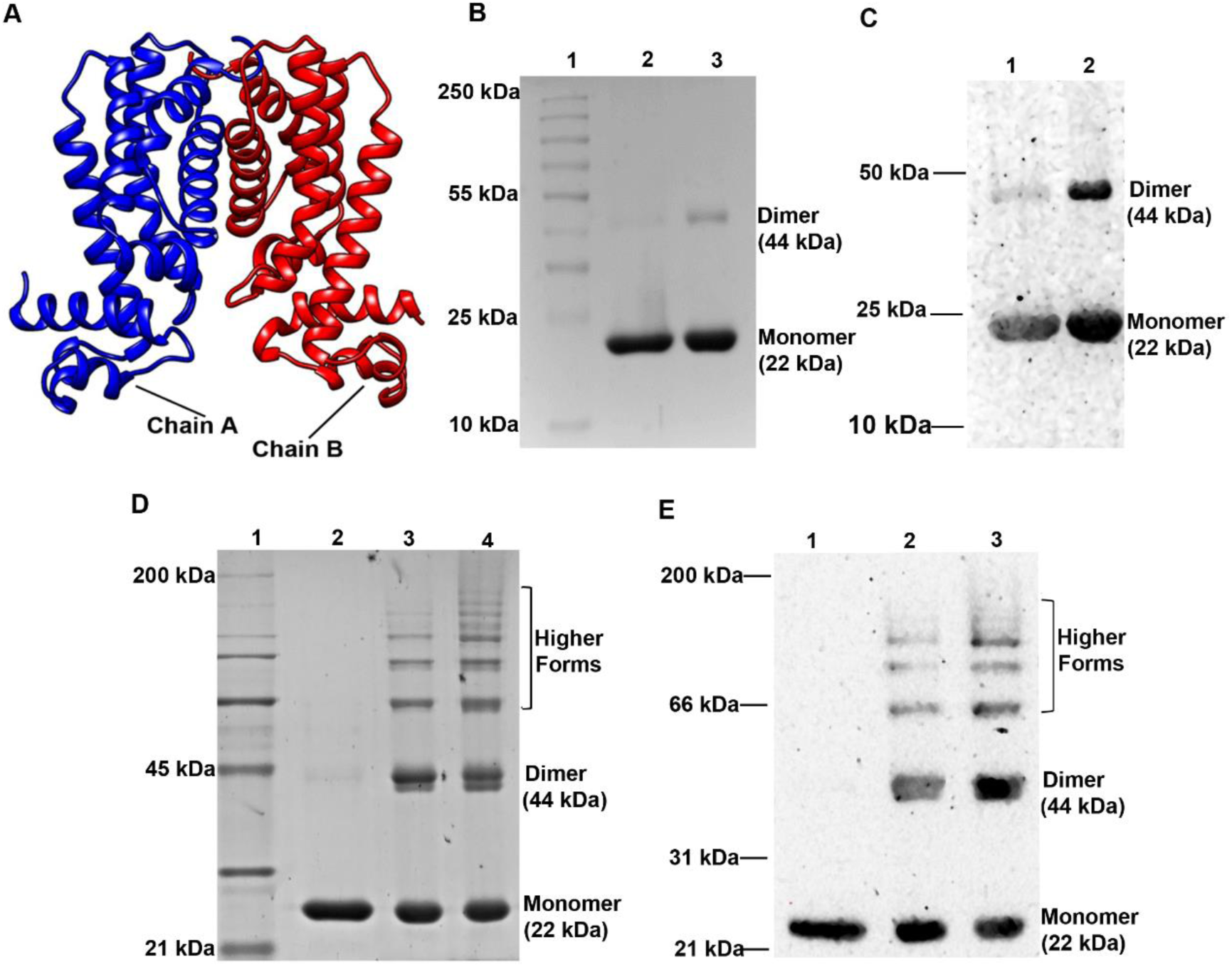
Rv1019 forms dimers. **(A)** *In silico* model shows dimerization of Rv1019 (homodimer –in blue and green); **(B)** Coomassie Brilliant Blue stained of polyacrylamide gel shows dimer formation *in vitro* under non-denaturing condition. Lane 1: Protein ladder, Lane 2: Rv1019 treated with SDS-PAGE loading dye containing SDS, β-mercaptoethanol, DTT, and was heated (control), Lane 3: Rv1019 in loading dye without SDS, β-mercaptoethanol, DTT, and was not heated **(C)** Western blot using antibodies against Rv1019 to confirm the monomer and dimer forms of Rv1019. Lane 1: Rv1019 treated with loading dye containing SDS, β-mercaptoethanol, DTT, and was heated (control), Lane 2: Rv1019 treated with loading dye without SDS, β-mercaptoethanol, DTT and was not heated; **(D)** Rv1019 forms dimers and higher forms upon cross-linking with glutaraldehyde. Lane 1: Protein ladder, Lane 2: Rv1019 treated with loading dye containing SDS, β-mercaptoethanol, DTT, and was heated (control), Lanes 2-3: Rv1019 treated with 0.0001% and 0.0005% glutaraldehyde, respectively, and loaded with loading dye without SDS, β-mercaptoethanol, DTT and was not heated; **(E)** Western blot using antibodies against Rv1019 to confirm dimer and higher forms of Rv1019. Lane 1: Rv1019 treated with SDS-PAGE loading dye containing SDS, β-mercaptoethanol, DTT and heated (control), Lanes 2-3: Rv1019 treated with 0.0001% and 0.0005% glutaraldehyde, respectively, and loaded with loading dye without SDS, β-mercaptoethanol, DTT and was not heated.

### Binding of Rv1019 to the promoter is abrogated in the presence of tetracycline

Rv1019 belongs to TetR family of transcriptional regulators which often bind to DNA with an N-terminal DNA binding region containing helix-turn-helix domain. Tetracycline is known to bind to the C-terminal of these proteins which would cause a conformational change leading to the abrogation of its DNA binding activity. We generated the structure of Rv1019 by homology modelling using SWISS-Model server, and subjected it to docking analysis on PatchDock server. Docking of Rv1019 with tetracycline as ligand showed that tetracycline can interact with the protein (Fig. 4A). Further, addition of tetracycline (100 ng/ml) to *M. smegmatis* expressing Rv1019 under its own promoter showed significant increase in GFP expression (Fig. 4B and **Fig. S5**). To examine the effect of tetracycline on DNA binding, we performed an EMSA on ^32^P-labeled 18 bp palindrome with 10 μM of Rv1019 and increasing concentrations of tetracycline (1-5 μg). DNA-protein complex formation was found to decrease with increasing the concentrations of tetracycline suggesting the classical de-repression of a TetR family member (Fig. 4C).

**Fig. 4.**
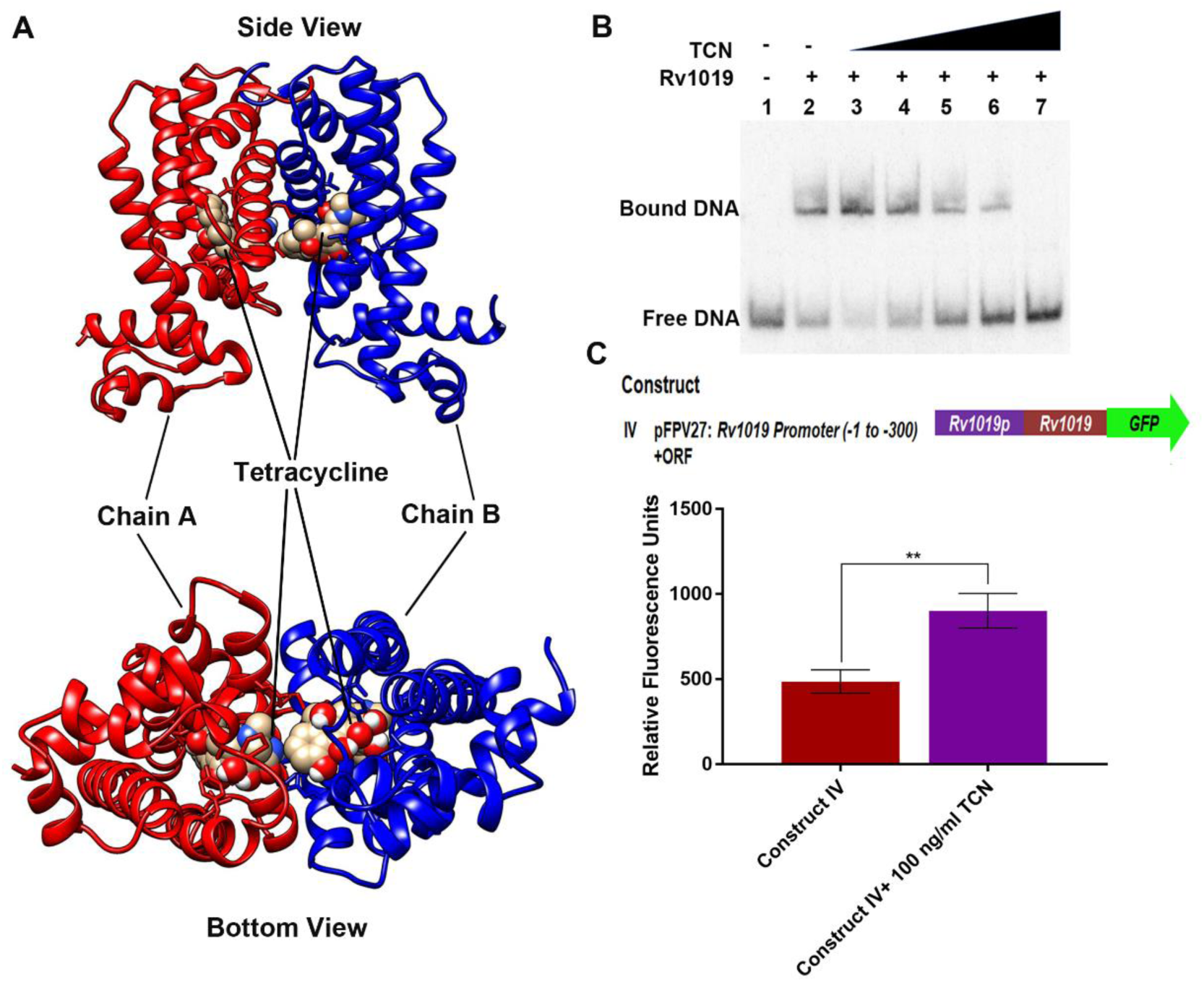
Tetracycline abrogates binding of Rv1019 to DNA. **(A)** Docking analysis shows tetracycline bound to Rv1019. **(B)** EMSA shows tetracycline abrogates DNA-Rv1019 interaction. Lane 1: ^32^P-labelled 18 bp inverted repeats in the absence of Rv1019, Lane 2: ^32^P-labelled DNA incubated with 10 of µm Rv1019 (positive control), Lanes 3-7: EMSA in the presence of increasing concentrations (1-5 µg) of tetracycline. **(C)** GFP reporter assay in the presence of tetracycline. Log phase *M. smegmatis ΔMSMEG_5424* harboring construct IV was treated with tetracycline (100 ng/ml) and fluorescence was measured after 3 h and represented in terms of RFU. GFP expression in *M. smegmatis* carrying construct IV vs construct IV in the presence of tetracycline was compared and values are represented as the mean ± standard deviation of three independent experiments (Student’s *t*-test). \*\*\**P* ≤ 0.005

### *Rv1019*, *Rv1020* and *Rv1021* are co-transcribed

Homologues of *Rv1019* locus are conserved across mycobacterium species with homologues of genes *Rv1020* and *Rv1021* downstream (Fig. 5A). *Rv1020* (*mfd*) codes for a transcription-coupled repair protein also known as transcription-repair coupling factor (TRCF), and *Rv1021* (*mazG*) codes for a nucleotide triphosphate phosphohydrolase. To check whether these three genes are transcribed together, we designed PCR primers that can amplify the junctions between the genes (Fig. 5B). By performing PCR using cDNA of the normally grown *Mtb* we were able to amplify the *Rv1019-Rv1020*, *Rv1020-Rv1021* and *Rv1019-Rv1021* junctions (Fig. 5C **and D)**. These results indicate that *Rv1019*, *Rv1020*, *Rv1021* are transcribed together into a single mRNA.

**Fig. 5.**
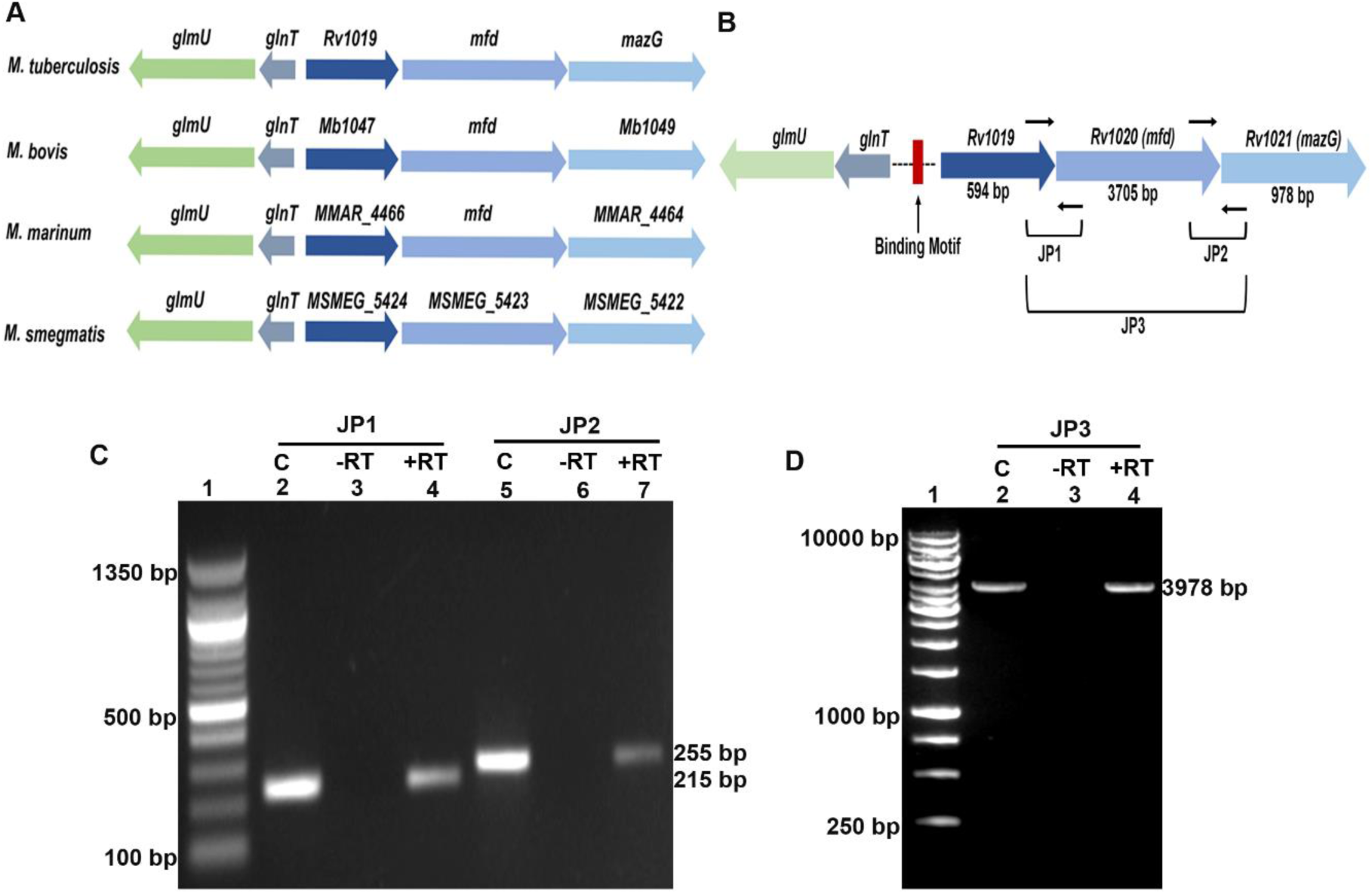
Rv1019, Rv1020 and Rv1021 are co-transcribed. **(A)** Representative gene maps of Rv1019, its homologues and gene neighbourhood in different *Mycobacterium* species; **(B)** Representative image shows the positions of the PCR primers designed to validate the co-transcription. Thin black arrows represent the DNA region selected to design PCR primers; **(C)** PCR amplification to prove co-transcription of *Rv1019, Rv1020, Rv1021* genes. JP1 (Junction Primer set 1-used to amplify *Rv1019-1020*), JP2 (Junction Primer set 2-used to amplify *Rv1020-1021*). Lane 1: 100 bp DNA ladder, Lanes 2 and 5: positive control, Lanes 3 and 6: PCR amplification from negative reverse transcription reaction, Lanes 4 and 7: PCR amplification from positive reverse transcription reaction; **(D)** PCR showing amplification of the entire transcript. JP3 (Junction Primer set 3-used to amplify *Rv1019-1021*). Lane 1: 1kb DNA ladder, Lanes 2: positive control, Lane 3: PCR amplicon from negative reverse transcription reaction, Lane 4: PCR amplification from of positive reverse transcription reaction.

### Rv1019 negatively regulates *Rv1020* and *Rv1021* expression

Gene map shows that *Rv1019* and its gene neighbourhoods are conserved in the surrogate host *M. smegmatis* (Fig. 6A). Virtual footprint analysis showed that the cognate binding site of Rv1019 that we identified in *Mtb* is also conserved in *M. smegmatis* with two mismatches (Fig. 6B). Therefore, we analysed whether Rv1019 is able to bind to the *M. smegmatis* regulatory sequence. EMSA with this template showed formation of DNA-protein complex as a function of increasing concentrations of recombinant Rv1019 (2-10 µM) (Fig. 6C). Specificity of binding was further confirmed by competitive and non-competitive EMSA (Fig. 6D).

**Fig. 6.**
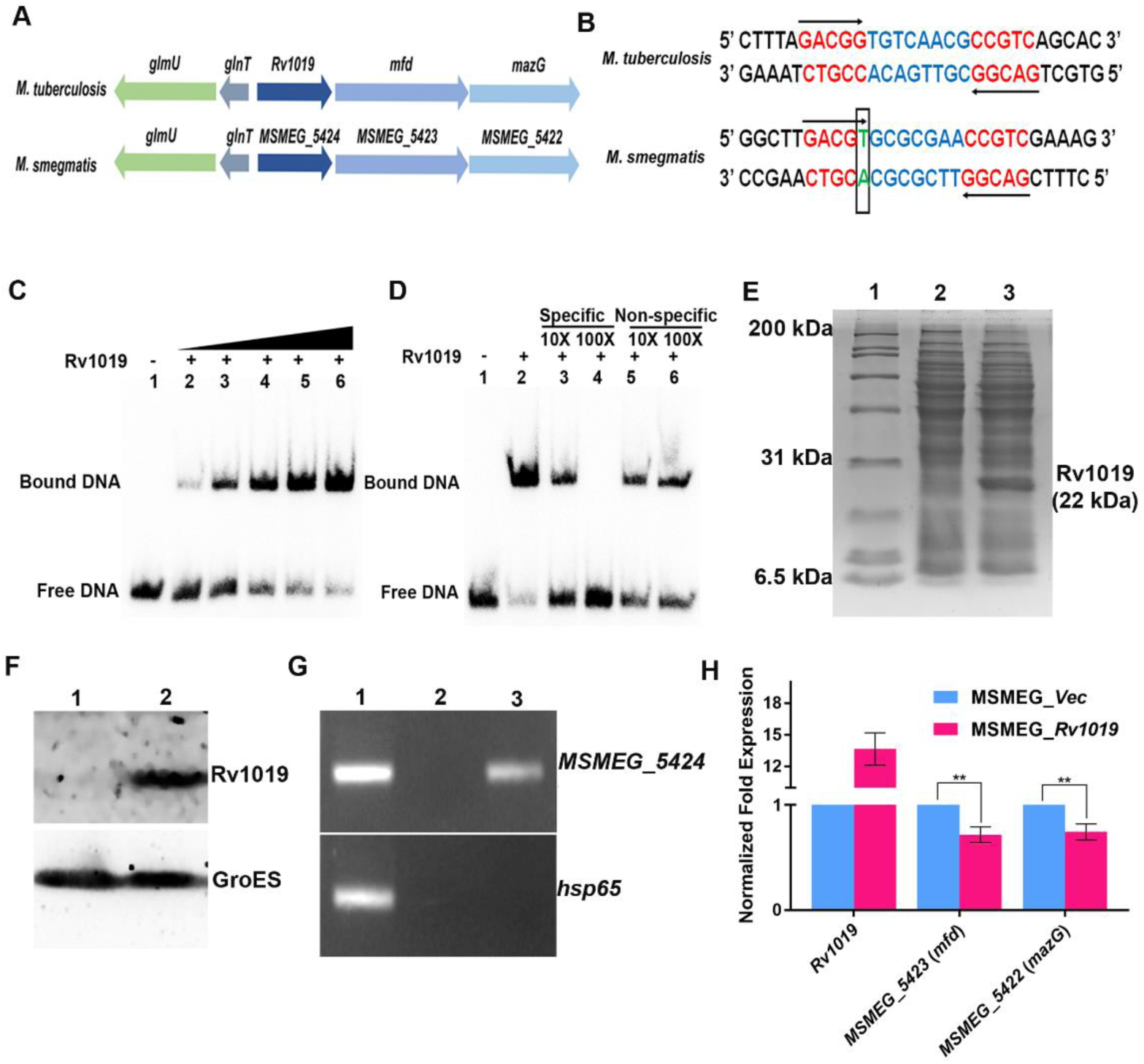
Rv1019 negatively regulates expression of *mfd* and *mazG*. **(A)** Gene map of *Rv1019*, *Rv1020* and *Rv1021* and their homologues in *M. smegmatis MSMEG_5424* (*Rv1019*), *MSMEG_5423* (*Rv1020* or *mfd*) and *MSMEG_5422* (*Rv1021* or *mazG*); **(B)**17 bp inverted repeats in *M. smegmatis* with one mismatch (green) and 7 bp gap region (blue); **(C)** EMSA shows binding of Rv1019 to the cognate binding region in the *M. smegmatis* DNA. Lane 1: 17 bp palindrome DNA without protein, Lanes 2-6 increasing concentrations of Rv1019 (2-10 µM); **(D)** Competitive and non-competitive EMSA to validate specificity of binding. Lane 1: ^32^P-labeled DNA without protein, Lane 2: ^32^P-labelled DNA incubated with 10 µm of Rv1019 (positive control), Lanes 3-4: competitive EMSA, Lanes 5-6: non-competitive EMSA; **(E)** *Rv1019* constitutive expression in *M. smegmatis.* Lane 1: Protein ladder, Lane 2: Vector control (*M. smegmatis: pBEN*), Lane 3: *M. smegmatis: pBEN:Rv1019*; **(F)** Western blot using antibodies against Rv1019 to confirm *Rv1019* expression. Lane 1: Vector control (*M. smegmatis*: *pBEN*), Lane 2: *M. smegmatis: pBEN: Rv1019*; **(G)** ChIP assay showing binding of Rv1019 to cognate binding region in *M. smegmatis* DNA. Lane 1: input DNA, Lane 2: pulldown by IgG antibody (negative control), Lane 3: pulldown with Rv1019 antibodies; **(H)** Quantitative real-time PCR showing expression of *Rv1019*, *MSMEG_5423* and *MSMEG_5422* in *M. smegmatis* expressing *Rv1019*. Values are the mean ± standard deviation of three independent experiments (Student’s *t*-test). **P ≤ 0.002

In order to check the hypothesis whether the auto-repressor activity of Rv1019 affects the expression of downstream genes, we constitutively expressed *Rv1019* in *Mtb*. Expression was confirmed by SDS-PAGE and western blot (Fig. 6E and **F)**. *In vivo* binding of Rv1019 on *M. smegmatis* DNA sequence was confirmed by ChIP using Rv1019 antibodies (Fig. 6G). Expression of *MSMEG_5424* (*Rv1019)*, *MSMEG_5423 (Rv1020)* and *MSMEG_5422 (Rv1021)* was analysed by qRT-PCR in *M. smegmatis* expressing *Rv1019*. We found that the Rv1019 constitutively expressing strain showed significant reduction of expression of *Rv1020* and *Rv1021* compared to the vector control (Fig. 6H).

### *M. smegmatis* expressing *Rv1019* is sensitive to DNA damaging agents

Mfd and MazG play a critical role in DNA repair in *Mtb* (22). To check whether Rv1019 has any role in DNA repair we exposed *M. smegmatis* expressing *Rv1019* to UV light and H_2_O_2_, which cause DNA lesions in bacteria. *M. smegmatis* expressing *Rv1019* showed increased sensitivity to UV light compared to the vector control strain. *M. smegmatis* lacking *recA* served as a control (Fig. 7A and 7B). At the same time at 5mM H_2_O_2_ *M. smegmatis* expressing *Rv1019* showed increased sensitivity to oxidative stress compared to the vector control strain (Fig. 7C and 7D). These results show that *M. smegmatis* expressing *Rv1019* is more sensitive to DNA damaging agents.

**Fig. 7.**
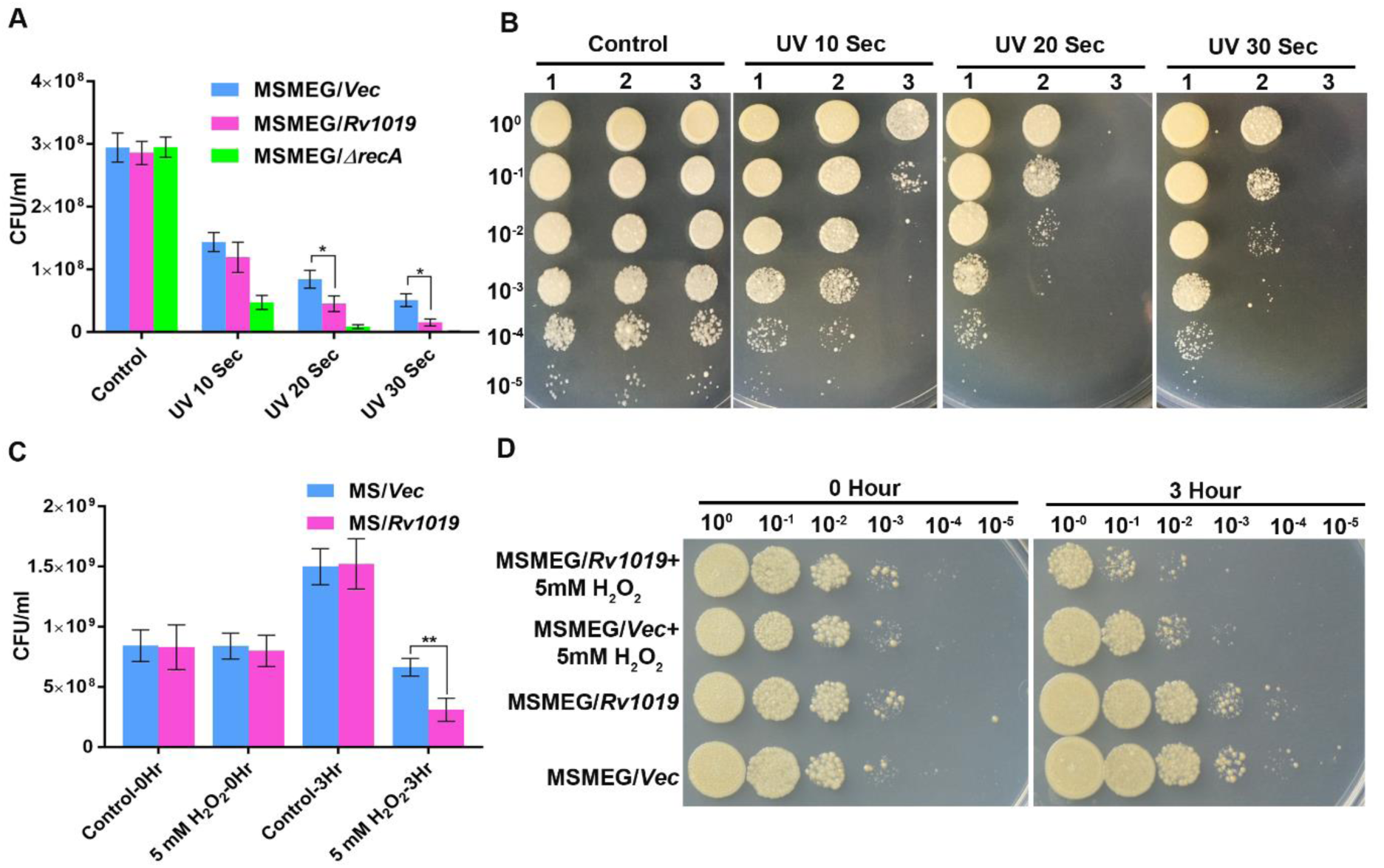
*M. smegmatis* expressing *Rv1019* is sensitive to UV light and H_2_O_2_. **(A)** Viable count of *M. smegmatis* after exposure to UV radiation at various time points. Colonies were counted and *M. smegmatis* harbouring empty vector was compared with bacteria expressing *Rv1019* and the values are the mean ± standard deviation of three independent experiments (Student’s *t*-test). *P ≤ 0.01; **(B)** Spot assay. Unexposed bacterial cultures and those exposed to UV light were serially diluted and spotted on Middlebrook 7H10 agar plates. Colonies were observed after 48 h of incubation. Lane 1: *M. smegmatis:pBEN*, Lane 2: *M. smegmatis:pBEN:Rv1019,* Lane 3: *M. smegmatis:ΔrecA*; **(C)** Hydrogen peroxide sensitivity assay. Viable count of *M. smegmatis* after treatment with 5mM H_2_O_2_ for 3 h. Colonies were counted and *M. smegmatis* with empty vector was compared with bacteria expressing *Rv1019,* and the values are the mean ± standard deviation of three independent experiments (Student’s *t*-test) **P ≤ 0.008; **(D)** Spot assay. Untreated cells and those treated with H_2_O_2_ were serially diluted and spotted on 7H10 agar plates. Colonies were observed after 48 h of incubation at 37 °C.

## Discussion

Success of *Mtb* as an intracellular pathogen depends on its ability to overcome the challenging environment in the host, and the bacterium still continues to be the leading cause of death by an infectious agent. In the host the bacteria enter a metabolically inactive, non-replicating dormant phase called latent TB infection (LTBI), and later gets reactivated under immune-compromised conditions. Transcriptional regulation in intracellular pathogens plays an important role in adapting to the challenging environment inside the host cells. Some of the key transcriptional regulators expressed in *Mtb* during persistence inside the human body include, DevR, KstR, etc. The focus of our study was to characterize Rv1019, a putative transcriptional regulator of *Mtb* H37Rv, that was found to be differentially expressed in *in vitro* dormancy-reactivation models (19, 20).

Using *M. smegmatis*, the surrogate organism that is widely used to study the biology of *Mtb*, we analysed the auto-regulatory mechanisms of Rv1019 *in vivo*. With the help of a series of reporter constructs we showed that Rv1019 is an auto-repressor. By employing *in vitro* DNA-protein interaction assays and online bio-informatics tools we showed that it binds to the promoter of its own gene, and the cognate binding site is −146 to −129 bp region upstream of its ORF. Binding of Rv1019 to an incomplete inverted repeat in its promoter showed the typical binding preference of a transcriptional regulator of TetR family (7).

Formation of a homodimer or multimer is a key feature of this class of proteins which favours DNA binding (8, 23), and we demonstrated that Rv1019 can form dimers *in vitro*. *In silico* studies showed tetracycline can bind to the dimer altering its DNA binding ability. DNA-protein interaction assay also showed the abrogation of binding in the presence of tetracycline confirming that Rv1019 is a classical transcriptional regulator of TetR family which could be influenced by the binding of a ligand.

Many of the TetR family transcriptional regulators are found to be regulating the expression of the gene neighbourhood. For example, transcriptional regulator *Rv1219c* represses the expression of *Rv1217c* and *Rv1218c* involved in multidrug efflux transport system (23, 24). Information provided by the mycobacterium databases shows that two genes, *Rv1020* and *Rv1021*, are present in the same orientation downstream of *Rv1019*. *Rv1020* codes for Mfd, also known as transcription-repair coupling factor (TCRF), and *Rv1021* codes for MazG, a nucleoside triphosphate pyrophosphohydrolase, which are two key proteins involved in DNA repair in *Mtb* (22, 25). This prompted us to investigate the effect of Rv1019 on the expression of these downstream genes. By RT PCR analysis of gene junctions in the mRNA we showed that *Rv1019*, *Rv1020* and *Rv1021* are transcribed into a single mRNA molecule. Over-expression of *Rv1019* significantly reduced the expression of orthologues of *mfd* and *mazG* in *M. smegmatis* showing that Rv1019 is a negative regulator of these downstream genes. Therefore, we propose that these three genes are in a single operon and *Rv1019* is the first gene of *Rv1020*-*Rv1021* operon and acts as the key regulator. Our results correlate with the existing transcriptomic and operon databases of mycobacterium where it is predicted to be co-transcribed with *mfd* and *mazG* (26–28). *Rv1019*, *Rv1020*, *Rv1021* gene locus is conserved in pathogenic mycobacteria such as *M. tuberculosis*, *M. leprae*, *M. bovis* and *M. marinum* as well as in non-pathogenic *M. smegmatis*. Mycobacterium is equipped with transcription-coupled repair (TCR), an effective method of detection and repair of DNA lesions caused by DNA damaging agents such as UV radiation and various oxidative stresses (29). UvrA, UvrB, UvrC, UvrD and Mfd of TCR system are essential for the repair of DNA lesions resulting from exposure to such DNA damaging environments (29–31). Our results showed increased sensitivity of *M. smegmatis* expressing *Rv1019* to UV light. In intracellular pathogens such as *Mtb* UV radiation is unlikely to induce activation of TCR system, although DNA-damaging factors such as oxidative stress within the macrophages are likely to be possible inducers of the same. This assumption was supported by the significant reduction in the number of *M. smegmatis* expressing *Rv1019,* when the bacterium was subjected to oxidative stress. However, at this point we are unable to affirm whether this susceptibility is driven by the downregulation of *Mfd* or through an *mfd*-independent mechanism induced by Rv1019. A recent report suggests that Mfd could act as an ‘evolvability factor’ promoting antimicrobial resistance in *Mtb* (29). It will be interesting to know if Rv1019 plays the role of a key regulator of genetic changes in *Mtb* leading to drug resistance.

Mycobacterial MazG is involved in DNA synthesis by the hydrolysis of dNTPs and elimination of mutagenic dNTPs generated by oxidative stresses (32–34). Deletion of *mazG* has shown to cause genomic instability in dormant *Mtb* (34), whereas antibiotic-induced oxidative stress reduced the survival of *ΔmazG Mtb* in activated macrophages (33). We found that downregulation of the expression of *MSMEG_5422*, the orthologue of *mazG,* in *M. smegmatis* constitutively expressing *Rv1019,* made the bacteria more sensitive to H_2_O_2._ These observations suggest a role of this protein in oxidative stress response in *Mtb*.

Thus, in our effort to functionally characterize Rv1019, a putative TetR family transcriptional regulator of *Mtb*, we found a novel regulatory mechanism by which this protein regulates DNA damage response which is essential for the bacterium for its survival under the hostile conditions within the host and contributes to its persistence. The role of Rv1019 under dormancy-reactivation and different stress conditions, and its significance in the pathogenesis of *Mtb* needs further analysis. Transcriptomic, proteomic and DNA-protein interaction studies will reveal the regulatory networks and interactions which may provide better understanding of the significance of Rv1019 in the growth and survival of *Mtb*. As Rv1019 could regulate expression of *mfd* and *mazG* which code for two key proteins involved in DNA repair system of *Mtb*, we propose this regulator is a prospective target for therapeutic intervention to fight tuberculosis, especially latent TB infection. Inhibitors of such regulators of critical pathways that help *Mtb* to persist in human body could possibly be used as prophylactic drugs against TB.

## Materials and Methods

### Bacterial strains, plasmids and growth conditions

*Rv1019* was cloned and expressed in *E. coli* JM109 (Promega, Madison, WI, USA) and *E. coli* BL21 (DE3) (Novagen, Madison, WI, USA), respectively. All *E. coli* strains used in this study were grown in Luria Bertani (LB) broth or on LB agar (HiMedia, Mumbai, India) at 37 °C. *M. tuberculosis* and *M. smegmatis* were cultured in Middlebrook 7H9 (Becton Dickinson, Franklin Lakes, NJ, and USA) supplemented with 10% albumin-dextrose-catalase (ADC) HiMedia, Mumbai, India. All steps involving handling of *Mtb* were carried out in a biosafety level three (BSL3) facility. For cloning and over-expression of proteins in *E. coli,* pET-32a plasmid (Novagen) was used, for promoter analysis in *M. smegmatis* pFPV27 vector was used. For generating knockout of *MSMEG_5424*, pKM342 and pJV53 (Addgene) were used. PCR primers and oligos were purchased from Sigma-Aldrich. Restriction enzymes were purchased from New England Biolabs, Ipswich, MA, USA. All other chemicals and reagents were purchased from Sigma-Aldrich, St. Louis, MO, USA or USB, Cleveland, OH, USA.

### Cloning and expression of *Rv1019* in *E. coli*, and purification of the recombinant protein

*Rv1019* ORF was amplified from the genomic DNA of *Mtb* H37Rv using gene specific primers and cloned into pET-32a expression vector (**Table. S1** and **Table. S2**) with a 6xHis tag and transformed into *E. coli* BL21 DE3 cells. Protein expression was induced by adding 1 mM IPTG (isopropyl β-D-1-thiogalactopyranoside) and by incubating at 37° C for 8 h. Protein was purified by Ni-NTA chromatography and subjected to dialysis in 50mM NaH_2_PO4 and 150 Mm NaCl and concentrated using 3K centrifugal concentrator (Pall Corporation, Ann Arbor, MI, USA) and stored at −20 °C in 30% glycerol.

### Preparation of Antibodies

Recombinant Rv1019 was injected intravenously into male New Zealand White rabbits for raising polyclonal antibodies. Procedures for immunization was carried out at Rajiv Gandhi Centre for Biotechnology according to the guidelines of Institutional Animal Ethics Committee. Immunization, serum collection, antibody purification and western blot against *Mtb* cell lysate were carried out as described by Gomez et al (35). Briefly, purified Rv1019 protein (200μg) was emulsified in TitreMax Gold adjuvant (Sigma-Aldrich) and used to immunize two one-year old male rabbits by subcutaneous injection. Antiserum was collected at the fourth week and a booster immunization was given on the same day. Antibodies were purified using AminoLink Resin (Thermo Scientific) according to the manufacturer’s instructions. Purified antibodies were stored as aliquots supplemented with 0.02% sodium azide (Sigma-Aldrich) at −20 °C.

### Glutaraldehyde cross-linking of Rv1019

Cross-linking experiment was performed as describe by Gomez et al (35). Briefly, recombinant Rv1019 was treated with 0.0002 and 0.0005% of glutaraldehyde for 30 min at 37 °C. Cross-linking was terminated by adding PAGE sample loading dye with 25 mM DTT and resolved on a 12% polyacrylamide gel in the absence of SDS. Gel was stained with Coomassie Brilliant Blue G-250. Monomer and dimer bands were excised and subjected to MALDI-TOF/MS/MS analysis (ultrafleXtreme; Bruker Daltonik, Bremen, Germany). The data obtained were compared with that in NCBI non-redundant database using MASCOT (version 2.2.04; Matrix Science, London, UK).

### Electrophoretic mobility shift assay (EMSA)

All the templates required for the assay were labelled with [γ-^32^P] ATP using T4 DNA ligase (NEB) at room temperature for 1 h. Electrophoretic mobility shift assays were carried out in 20 µl reactions containing labelled template DNA (5 nM), increasing concentrations of Rv1019 (2-10 µM), 1X buffer containing 10 mM Tris pH 8.0, 1 mM EDTA pH 8.0, 1mM DTT, 10% glycerol and BSA (100 µg/ml). DNA fragments used for EMSA were prepared either by annealing complementary oligos or by PCR **(Table. S1)**. Increasing concentrations of tetracycline (1-5μg) were added to the reaction mixture to test its effect on the DNA binding property of Rv1019. Binding reactions were carried out for 30 min at 25 °C. Samples were electrophoresed on 5% polyacrylamide gel at 100 V at 25 °C. Gel was dried and exposed to Phosphor Imager (Bio-Rad) and scanned with Typhoon FLA 9500 (GE Life sciences, Pittsburgh, PA, USA).

### Reporter constructs and spectrofluorimetry

pFPV27, a promoter-less reporter plasmid was used to make the reporter constructs. To study the activity of the promoter of *Rv1019*, a 300 bp DNA fragment upstream from its start site was amplified from *Mtb* H37Rv genomic DNA, and cloned into pFPV27, at *Apa*I and *Eco*RI restriction sites upstream of GFP ORF to generate transcriptional fusions (**Table. S2**). To study the effect of Rv1019 on its own promoter, a reporter plasmid was constructed by cloning *Rv1019* along with the upstream 300 bp region in pFPV27. These constructs were electroporated into *M. smegmatis* using Gene Pulser Xcell (Bio-Rad) and the transformants were selected on 7H10 agar plates containing kanamycin (50 μg/ml). The recombinant *M. smegmatis* was grown in LB medium and the fluorescence was measured after 48 h. Optical density was measured at 600 nm and fluorescence was measured at the emission maximum of 520 nm using an excitation of 488 nm in a microplate reader (Tecan, Mannedorf, Switzerland). Relative fluorescence units (RFU) were calculated by dividing fluorescence units ^b^y *D*_600_ values. *M. smegmatis* harbouring empty pFPV27 vector and *M. smegmatis* harbourin^g^ ^p^FPV27 containing *hsp60* promoter were used as controls. All measurements were correcte^d^ for background fluorescence. The experiments were carried out in triplicate.

### RNA isolation, cDNA synthesis and quantitative PCR

RNA was isolated from *Mtb* and *M. smegmatis* using the protocol described by Gopinath et al (19). Briefly, RNA was extracted using Trizol (Sigma-Aldrich) reagent and precipitated with 100% isopropanol. RNA was resuspended in nuclease-free water and treated with DNaseI (Sigma-Aldrich). Complementary DNA (cDNA) was prepared using Reverse Transcriptase Core kit (Promega Corporation) according to the manufacturer’s protocol. A qPCR was performed using SYBR Green kit (Takara Bio INC). The thermal cycling protocol was set as follows: an initial denaturation of 95 °C for 1 min; 35 cycles of denaturation at 95 °C for 30 s, primer annealing at 60 °C for 45 s, and extension at 72 °C for 30 s. Following amplification, a melt curve analysis was performed to confirm the specificity of the amplified product. Relative changes in gene expression were calculated using the −ΔΔ*C*_t_ method (36), with sigma factor A (*sigA*) as the housekeeping control gene.

### Chromatin immunoprecipitation assay

Chromatin immunoprecipitation (ChIP) assay was performed according to the protocol by Kahramanoglou et al (37). Briefly 20 ml of mid-log phase *Mtb* culture (OD_600_ ∼ 0.6) was treated with formaldehyde (1%), and incubated at 25 °C for 30 min to carry out DNA-protein cross-linking, and was terminated by addition of glycine to a final concentration of 0.5 M and incubating for 10 min at 25 °C. Cells were washed by centrifugation twice with 1× PBS (pH 7.4) containing 1 mM PMSF at 3000 g for 10 min at 4 °C. Pellets were then resuspended in 1ml of ChIP lysis buffer containing 50 mM Hepes/KOH (pH 7.5), 150 mM NaCl, 1 mM EDTA, 1% Triton X-100, 0.1% (w/v) sodium deoxycholate, 0.1% SDS, 0.1 mg/ml RNaseA, 1× protease inhibitor cocktail (Sigma-Aldrich) and 1 mM PMSF. Cells were lysed using zirconium beads (0.1 mm) in a Mini Bead Beater (Biospec, Bartlesville, OK, USA) for three pulses, 30 s each at 4200 rpm, with 1 min intervals on ice. DNA was sheared using a Bioruptor sonicator (Diagenode, Seraing, Belgium) with 35 cycles of 30 s on/off pulses to yield an average fragment size of ∼ 500 bp. Sheared fragments were recovered by centrifugation at 12000 g for 10 min at 4 °C. One hundred microliters of this lysate was used as the input control.

### UV Sensitivity Assay

*M. smegmatis* expressing Rv1019 and vector control were generated by electroporation using Gene pulse Xcell (Bio-Rad) and the recombinants were selected on agar plates containing kanamycin (50 µg/ml). Isolated colonies were further grown on Middlebrook 7H9 (Becton Dickinson, Franklin Lakes, NJ, and USA) on a 37 °C shaker incubator for 48 h. *M. smegmatis* lacking *recA* was used as control. An OD 0.5 of each strain was serially diluted and spotted onto Middlebrook 7H10 medium (Becton Dickinson, Franklin Lakes, NJ, and USA), exposed to UV light (25 mJ/cm^2^) for 10 sec, 20 sec and 30 sec. Exposed plates were incubated for 48 h in the dark in a 37 °C incubator. Plates that were not exposed to UV light served as control.

### Oxidative stress assay

*M. smegmatis* expressing *Rv1019* and vector control were grown to log phase in Middlebrook 7H9 medium (Becton Dickinson, Franklin Lakes, NJ, and USA) on a 37 °C shaker incubator. Optical density was adjusted to 0.5 and bacteria were treated with 5 mM H_2_O_2_. *M. smegmatis* carrying empty plasmid served as the control. Both strains were incubated on a 37 °C shaker incubator for 3 h post H_2_O_2_ treatment, and were serially diluted and plated as well as spotted on to Middlebrook 7H10 medium (Becton Dickinson, Franklin Lakes, NJ, and USA). Plates were incubated on a 37 °C incubator; colonies were counted and spots were observed after 48 h.

### Bio-informatics analysis

Mycobaterium database https://mycobrowser.epfl.ch/ was used to collect protein and DNA sequence information. DNA binding domain was analysed at https://prosite.expasy.org/. Palindromic sequences finder (http://www.biophp.org/minitools/find_palindromes/demo.php) was used to identify palindromes in the promoter region. Rv1019 binding sites in the whole *Mtb* genome were identified using Virtual Footprint (38). MEME was used to identify putative binding sites (39). MEME was set to find palindromic motifs with a minimum width of 6 bp and a maximum of 50 bp. MEME was set to return a maximum of three motifs. Analysis of dimer formation was carried out at SWISS-MODEL server (https://swissmodel.expasy.org/interactive) and docking studies were carried out in PatchDock server (https://bioinfo3d.cs.tau.ac.il/PatchDock/php.php) and analysed by UCSF Chimera version 1.13.1.

## Abbreviations

Mtb: *Mycobacterium tuberculosis*
TB: tuberculosis
TetR: tetracycline resistance regulator
ChIP: chromatin immunoprecipitation
EMSA: electrophoretic mobility shift assay
TCRF: transcription-repair coupling factor.

## Acknowledgement

A.R.P thanks the Department of Science and Technology, Government of India, for INSPIRE fellowship. R.A.K is grateful to the Department of Biotechnology, Government of India for financial support.

The funders had no role in study design, data collection and interpretation, or the decision to submit the work for publication.

We are grateful to Arumugam Rajavelu and Krishna Kurthkoti for their inputs.

A.R.P and R.A.K designed and conceived the experiments. R.A.K supervised the execution of the experiments and A.R.P performed the experiments. A.R.P and R.A.K wrote the manuscript.

## Supplementary Figure Legends

**Fig. S1 Cloning and expression of *Rv1019.* (A)** PCR amplification of *Rv1019* from genomic DNA of *Mtb* H37Rv using gene specific primers. Lane 1: 100 bp DNA ladder, Lane 2: *Rv1019* amplicon; **(B)** Expression of *Rv1019* in *E. coli* BL21 DE3 cells. Lane 1: Broad range protein standard, Lane 2: lysate of uninduced *E. coli* BL21 DE3 harbouring pET32-a:*Rv1019*, Lane 3: lysate of IPTG-induced *E.coli* BL21 DE3 harbouring pET32-a:*Rv1019*, Lane 4: purified recombinant Rv1019; **(C)** Western blot of recombinant histidine-tagged Rv1019 developed with Histidine tagged antibodies; **(D)** Western blot of Rv1019 developed with purified Rv1019 antibodies. Lane 1: purified recombinant Rv1019, Lane 2: *M. tuberculosis* H37Rv lysate.

**Fig. S2 Reporter assay in *M. smegmatis*. (A)** Generation of *M. smegmatis ΔMSMEG_5424*. Lane 1: 1kb DNA ladder, Lane 2: *MSMEG_5424* PCR amplicon, Lane 3: PCR amplicon of *MSMEG_5424* with upstream and downstream 600 bp region, Lane 4: PCR amplicon of *MSMEG_5424* in *M. smegmatis ΔMSMEG_5424*, Lane 5: PCR amplification of *MSMEG_5424* with upstream and downstream 600 bp region containing hygromycin; **(B)** PCR amplification of probable promoter region 300 bp upstream of *Rv1019*. Lane: 1: 100 bp DNA ladder, Lane 2: PCR amplicon of 300 bp upstream region, Lane 3: PCR amplicon of *Rv1019* ORF with probable promoter region of 300 bp. **(C)** Confocal images of reporter constructs I, II, III and IV. Scale bars are 5 µM.

**Fig. S3 Confocal images of *M. smegmatis* carrying reporter constructs.** GFP expression of constructs III, IV and VI. **(A)** PCR amplification of *Rv1019* ORF along with 100 bp upstream region lacking the cognate binding sites. Lane 1: 100 bp DNA ladder, Lane 2: PCR amplicon with *Rv1019* ORF with 100 bp upstream region (Construct VI); **(B)** Confocal images of *M. smegmatis* carrying constructs III, IV and VI. Scale bars are 5 µM.

**Fig. S4 MALDI-TOF/MS/MS analysis.** Rv1019 treated with glutaraldehyde was electrophoresed on a non-denaturing polyacrylamide gel and bands were excised and analysed by MALDI-TOF/MS/MS.

**Fig. S5 Confocal microscopy of *M. smegmatis* carrying Rv1019 promoter with ORF treated with tetracycline**. GFP expression in *M. smegmatis* carrying construct IV in the absence and presence of tetracycline. Scale bars are 5 µM.

**Supplementary Table 1 (Table. S1)-**Attached with supplementary information

**Supplementary Table 1 (Table. S2)-**Attached with supplementary information

