## Supplemental Material for "Putative TetR family transcriptional regulator Rv1019 of *Mycobacterium tuberculosis* is an auto-repressor and a negative regulator of *mfd-mazG* operon"

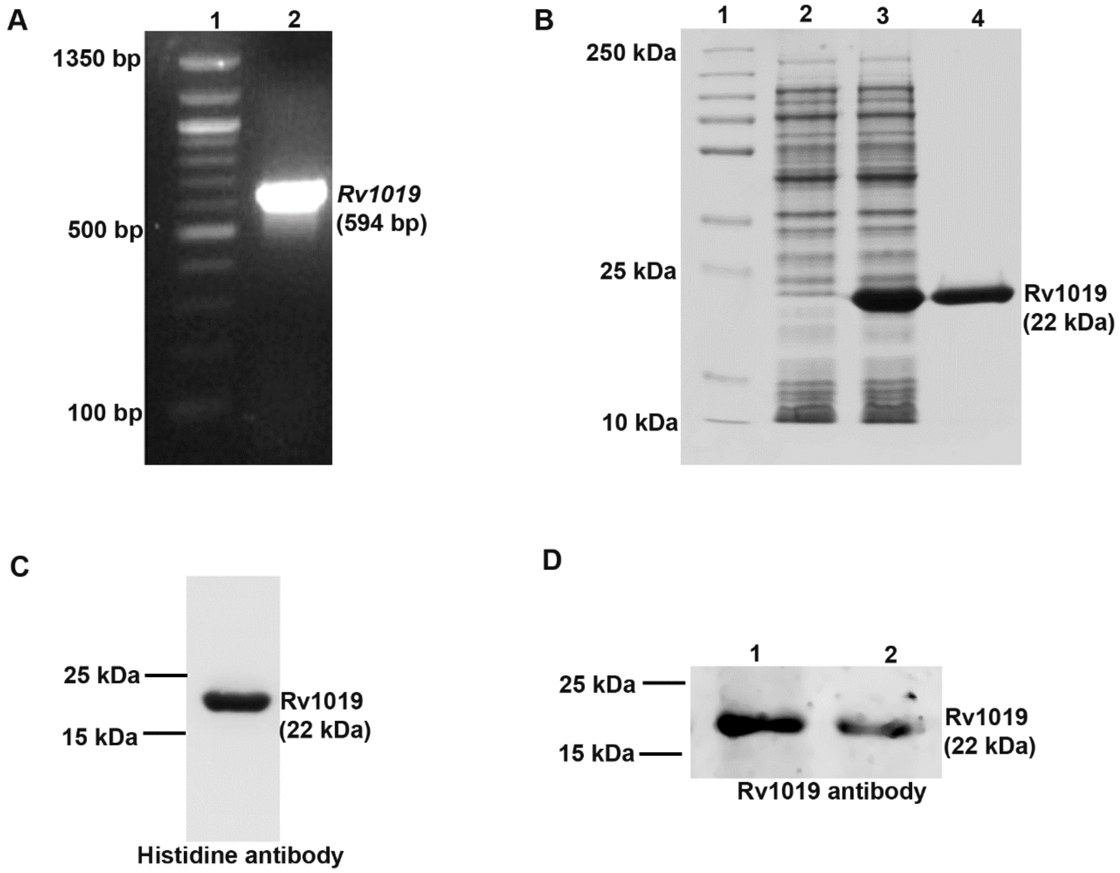


**Fig. S1 Cloning and expression of *Rv1019.* (A)** PCR amplification of *Rv1019* from genomic DNA of *Mtb* H37Rv using gene specific primers. Lane 1: 100 bp DNA ladder, Lane 2: *Rv1019* amplicon; **(B)** Expression of *Rv1019* in *E. coli* BL21 DE3 cells. Lane 1: Broad range protein standard, Lane 2: lysate of uninduced *E. coli* BL21 DE3 harbouring pET32-a:*Rv1019*, Lane 3: lysate of IPTG-induced *E.coli* BL21 DE3 harbouring pET32-a:*Rv1019*, Lane 4: purified recombinant Rv1019; **(C)** Western blot of recombinant histidine-tagged Rv1019 developed with Histidine tagged antibodies; **(D)** Western blot of Rv1019 developed with purified Rv1019 antibodies. Lane 1: purified recombinant Rv1019, Lane 2: *M. tuberculosis* H37Rv lysate.

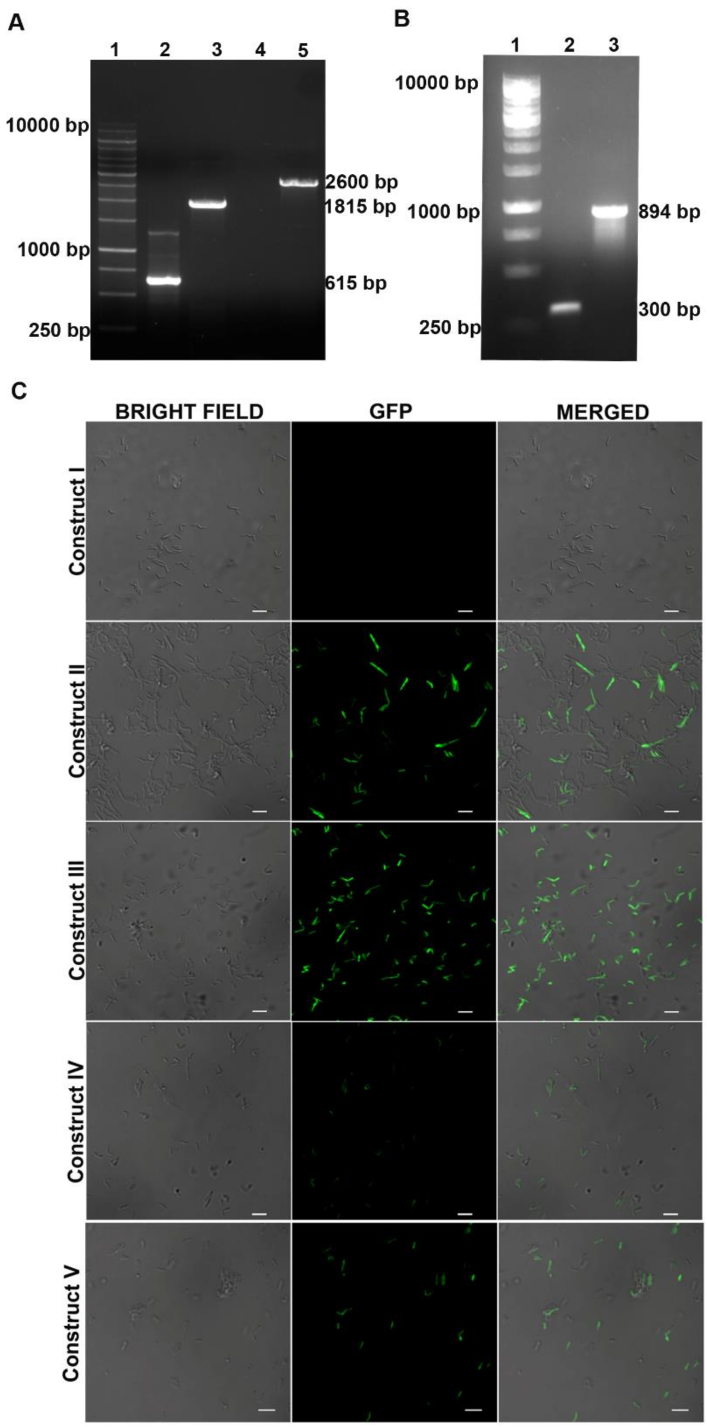


**Fig. S2 Reporter assay in *M. smegmatis*. (A)** Generation of *M. smegmatis ΔMSMEG_5424*. Lane 1: 1kb DNA ladder, Lane 2: *MSMEG_5424* PCR amplicon, Lane 3: PCR amplicon of *MSMEG_5424* with upstream and downstream 600 bp region, Lane 4: PCR amplicon of *MSMEG_5424* in *M. smegmatis ΔMSMEG_5424*, Lane 5: PCR amplification of *MSMEG_5424* with upstream and downstream 600 bp region containing hygromycin; **(B)** PCR amplification of probable promoter region 300 bp upstream of *Rv1019*. Lane: 1: 100 bp DNA ladder, Lane 2: PCR amplicon of 300 bp upstream region, Lane 3: PCR amplicon of *Rv1019*

ORF with probable promoter region of 300 bp. **(C)** Confocal images of reporter constructs I, II, III and IV. Scale bars are 5 μM.

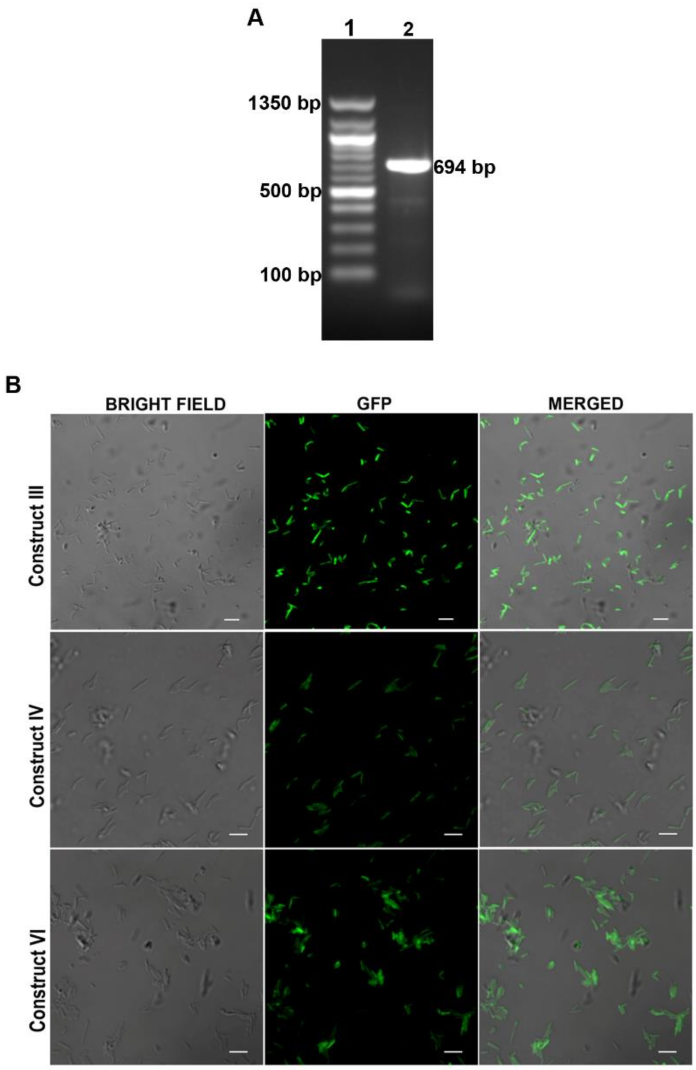


**Fig. S3 Confocal images of *M. smegmatis* carrying reporter constructs.** GFP expression of constructs III, IV and VI. **(A)** PCR amplification of *Rv1019* ORF along with 100 bp upstream region lacking the cognate binding sites. Lane 1: 100 bp DNA ladder, Lane 2: PCR amplicon with *Rv1019* ORF with 100 bp upstream region (Construct VI); **(B)** Confocal images of *M. smegmatis* carrying constructs III, IV and VI. Scale bars are 5 µM.

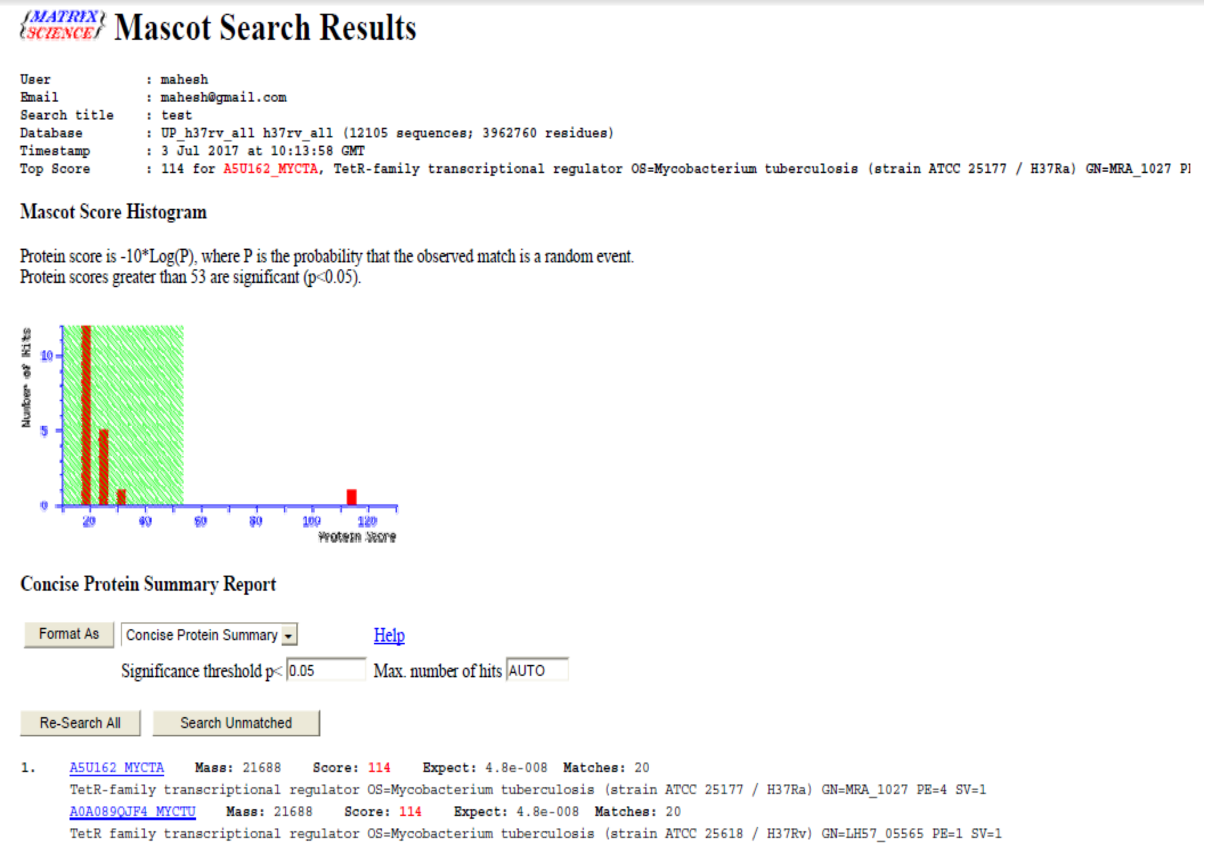

Mascot Search Results

User : mahesh

Search title : test
Database : UP_h37rv_all h37rv_all (12105 sequences; 3962760 residues)
Timestamp : 3 Jul 2017 at 10:13:58 GMT
Top Score : 114 for A5U162_MYCTA, TetR-family transcriptional regulator OS=Mycobacterium tuberculosis (strain ATCC 25177 / H37Ra) GN=MRA_1027 P[illegible]

Mascot Score Histogram

Protein score is -10*Log(P), where P is the probability that the observed match is a random event.
Protein scores greater than 53 are significant (p<0.05).



Concise Protein Summary Report

Format As | Concise Protein Summary | Help
Significance threshold p< 0.05 | Max. number of hits AUTO

Re-Search All | Search Unmatched

1. A5U162_MYCTA **Mass:** 21688 **Score:** 114 **Expect:** 4.8e-008 **Matches:** 20
TetR-family transcriptional regulator OS=Mycobacterium tuberculosis (strain ATCC 25177 / H37Ra) GN=MRA_1027 PE=4 SV=1
A0A089QJF4_MYCTU **Mass:** 21688 **Score:** 114 **Expect:** 4.8e-008 **Matches:** 20
TetR family transcriptional regulator OS=Mycobacterium tuberculosis (strain ATCC 25618 / H37Rv) GN=LH57_05565 PE=1 SV=1

**Fig. S4 MALDI-TOF/MS/MS analysis.** Rv1019 treated with glutaraldehyde was electrophoresed on a non-denaturing polyacrylamide gel and bands were excised and analysed by MALDI-TOF/MS/MS.

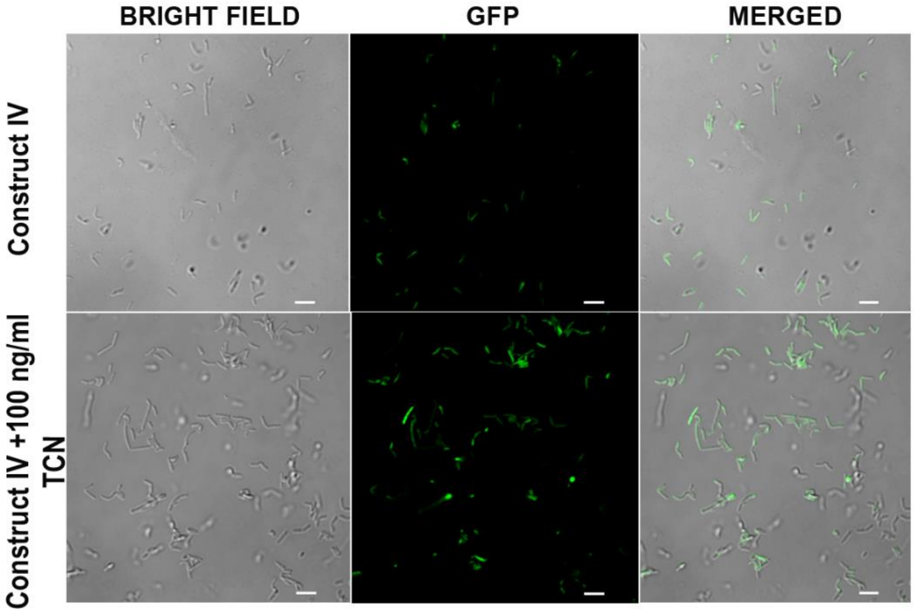


**Fig. S5 Confocal microscopy of *M. smegmatis* carrying Rv1019 promoter with ORF treated with tetracycline.** GFP expression in *M. smegmatis* carrying construct IV in the absence and presence of tetracycline. Scale bars are 5 µM.

**Table. S1: Bacterial strains and plasmids used in this study**

| **Strain or plasmid or primers** | **Description** | **Reference or source** |
|---|---|---|
| **Strains** | | |
| *E. coli* JM1019 | *traD36* *proA⁺B⁺ lacIq Δ(lacZ)M15/ Δ(lac-proAB) glnV44 e14- gyrA96 recA1 relA1 endA1 thi hsdR17* | Invitrogen, California, USA |
| *E. coli* BL21 DE3 | For inducible T7 expression; *E. coli* B $F^{-}$ *ompT gal [E. coli* B is naturally *dcm and lon] hsdS*$_B$ with DE3, a λ prophage carrying the T7 RNA polymerase gene and $lacI^Q$ | Invitrogen, California, USA |
| *Mycobacterium tuberculosis* H37Rv | Wild type | NIRT, Chennai, India |
| *Mycobacterium smegmatis* $mc^2155$ | Wild type | NIRT, Chennai, India |
| *Mycobacterium smegmatis* $mc^2155$: *ΔrecA* | *RecA* knock out strain of *Mycobacterium smegmatis* $mc^2155$ | Gift from Krishna Kurthkoti, RGCB, India |
| **Plasmids** | | |
| pET32-a | $Amp^r$; for production of recombinant protein | Invitrogen, California, USA |
| pKM342 | $Hyg^r$; allelic exchange vector | Gift from Krishna Kurthkoti |
| pFPV27 | $Kan^r$; promoter-less reporter system | Gift from Lalitha Ramakrishnan, Currently at University of Cambridge, UK |
| pFPV60 | $Kan^r$; *Hsp60* promoter reporter system | Gift from Lalitha Ramakrishnan |
| pBEN | $Kan^r$; *Hsp60* promoter mycobacterial shuttle vector | Gift from Lalitha Ramakrishnan |

**Table. S2: Primers and Oligos used in this study**

| Primer / Oligo Name | Sequence (5’ --------------- 3’) | Description |
|---|---|---|
| pET-*Rv1019* F | GGAATTCcatatgcatcaccatcaccatcacATGACGGGGACCGAGCGC | *Rv1019* amplification primer to clone into pET32-a, 6x-histifine tag (lower case underlined) forward, *Nde*I site (lower case) |
| pET-*Rv1019* R | CCCaagcttCTACTCGTCCTGTAGCCGCGG | *Rv1019* amplification primer to clone into pET32-a, reverse, *Hind*III site (lower case) |
| pKM-*MSMEG_5424* UP F | GGAATTCcatatgGACCGGTGAGCGTCGCGGT | *MSMEG_5424* upstream 500 bp amplification primer to clone into pKM342, forward, *Nde*I site (lower case) |
| pKM-*MSMEG_5424* UP R | GGAagatctTTCCTTGTCGGGTGCTGCCAC | *MSMEG_5424* upstream 500 bp amplification primer to clone into pKM342, reverse, *Bgl*II site (lower case) |
| pKM-*MSMEG_5424* DN F | CCGctcgagCCAGCGCGGGGTGGGTCAC | *MSMEG_5424* downstream 500 bp amplification primer to clone into pKM342, forward, *Xho*I site (lower case) |
| pKM-*MSMEG_5424* DN R | CCCaagcttCTTGCCGACCATGTCGACGCG | *MSMEG_5424* downstream 500 bp amplification primer to clone into pKM342, reverse, *Hind*III site (lower case) |
| pFPV-*Rv1019* Pro F | CGCggatccTTGGTTCAGATAGTTCAGGTTCG | *Rv1019* 300 bp upstream promoter amplification primer to clone into pFPV27, forward, *Bam*HI site (lower case) |
| pFPV-*Rv1019* Pro R | GATCggtaccCCTGGCGCGCGTCACCCG | *Rv1019* 300 bp upstream promoter amplification primer to clone into pFPV27, reverse, *Kpn*I site (lower case) |
| pFPV-*Rv1019* Pro+ ORF F | CGCggatccTTGGTTCAGATAGTTCAGGTTCG | Rv1019 300 bp upstream promoter with ORF amplification primer, forward, *Bam*HI site (lower case) |
| pFPV27-*Rv1019* Pro+ ORF R | GATCggtaccCTACTCGTCCTGTAGCCGCGG | Rv1019 300 bp upstream promoter with ORF amplification primer, reverse, *Kpn*I site ((lower case) |
| pFPV-*Rv1019* 100 bp pro + ORF F | CGCggatcc CACCGGCCCCACGCAAACC | *Rv1019* 100 bp upstream promoter without binding site + ORF amplification primer, forward, *Bam*HI site (lower case) |
| pFPV-*Rv1019* 100 bp pro + ORF F | GATCggtaccCTACTCGTCCTGTAGCCGCGG | *Rv1019* 100 bp upstream promoter without binding site + ORF amplification primer, reverse, *Kpn*I site (lower case) |

| | | |
|---|---|---|
| pBEN-*Rv1019* F | CGCggatccATGACGGGACCGAGCGC | *Rv1019* amplification primer to clone into pBEN, forward, *Bam*HI site (lower case) |
| BEN-*Rv1019* R | CCCAAGCTTCTACTCGTCCTGTAGCCGCGG | *Rv1019* amplification primer to clone into pBEN, reverse, *Hind*III site (lower case) |
| FR I F | GCCCGGCCGCTAGGACCA | EMSA fragment I primer, *Rv1019* upstream 300 bp, forward |
| FR I R | CCTGGCGCGCGTCACCCG | EMSA fragment I primer, *Rv1019* upstream -300 to -1 bp, reverse |
| FR II F | GCCCGGCCGCTAGGACCA | EMSA fragment II primer, *Rv1019* upstream -300 to -200 bp, forward |
| FR II R | GGTAGACGGCGACAGCGTTG | EMSA fragment II primer, *Rv1019* upstream -300 to -200 bp, reverse |
| FR III F | AATCAGAGGTGCTGCCGATTAC | EMSA fragment III primer, *Rv1019* upstream -200 to -100 bp, forward |
| FR III R | GGTAGACGGCGACAGCGTTG | EMSA fragment III primer, *Rv1019* upstream -200 to -100 bp, reverse |
| FR IV F | CACCGGCCCCACGCAAACC | EMSA fragment IV primer, *Rv1019* upstream -100 to -1 bp, forward |
| FR IV R | CCTGGCGCGCGTCACCCG | EMSA fragment IV primer, *Rv1019* upstream -100 to -1 bp, reverse |
| *Hsp60* Pro F | CGGTCATGGGCCGAACATAC | *Hsp60* promoter 300 bp upstream amplification primer, forward |
| *Hsp60* Pro R | TGCGAAGTGATTCCTCCGGATCGG | *Hsp60* promoter 300 bp upstream amplification primer, reverse |
| Palindrome Oligo S1 | TGActttagacggtgtcaacgccgtcagcacAGT | Rv1019 cognitive binding sequence (in lowercase), strand 1 |
| Palindrome Oligo S2 | ACTgaaatctgccacagttgcggcagtcgtgTCA | Rv1019 cognitive binding sequence (in lowercase), strand 2 |
| Random oligo S1 | GATCGATCGATCGATCGA | Non-specific binding sequence from *Hsp60* promoter, strand 1 |
| Random oligo S2 | CTAGCTAGCTAGCTAGCT | Non-specific binding sequence from *Hsp60* promoter, strand 2 |
| T4g32 S1 | GGGACCCTAGAGGTCCCCTTT | Transcription terminator sequence, strand 1 |
| T4g32 S2 | AAAGGGGACCTCTAGGGTCCC | Transcription terminator sequence, strand 2 |
| JP1 F | TGGTCAACCTGGTCTGGAATGG | Junction primer to amplify *Rv1019-Rv1020* junction, forward |
| JP1 R | TGCATGAGCTGTTGGAATGTCG | Junction primer to amplify *Rv1019-Rv1020* junction, reverse |

| | | |
|---|---|---|
| JP2 F | GCGCTCGCTGGGAAACCGC | Junction primer to amplify *Rv1020-Rv1021* junction, forward |
| JP2 R | CGTTTCCGGCGTGCGCCGA | Junction primer to amplify *Rv1020-Rv1021* junction, reverse |
| JP3 F | TGGTCAACCTGGTCTGGAATGG | Junction primer to amplify *Rv1019-Rv1021* junction, forward |
| JP3 R | CGTTTCCGGCGTGCGCCGA | Junction primer to amplify *Rv1019-Rv1021* junction, reverse |
| MS Pal S1 | GGCTTGACGTGCGCGAACCGTCGAAAG | Cognitive binding site in *M. smegmatis*, strand 1 |
| MS Pal S2 | CTTTCGACGGTTCGCGCACGTCAAGCC | Cognitive binding site in *M. smegmatis*, strand 2 |
| *Sig A* RT F | AGGATCAACTGCAGTCGGTG | qRT-PCR primer for sigma factor A, forward |
| *Sig A* RT R | CTTGGATTCGATCTGGCGGA | qRT-PCR primer for sigma factor A, reverse |
| *Rv1019* RT F | ATTCTGGCTGGAGACTTCGC | qRT-PCR primer for *Rv1019*, forward |
| *Rv1019* RT R | GCCATTCCAGACCAGGTTGA | qRT-PCR primer for *Rv1019*, reverse |
| *MSMEG_5423* (*mfd*) RT F | AGATGTCGACGATCCTCACC | qRT-PCR primer for *MSMEG_5423*, forward |
| *MSMEG_5423* (*mfd*) RT R | GGTTGTGGATGTAGAACGCC | qRT-PCR primer for *MSMEG_5423*, reverse |
| *MSMEG_5422* (*mazG*) qPCR F | CGCAGGAGAAGAAGGTCAAG | qRT-PCR primer for *MSMEG_5422*, forward |
| *MSMEG_5422* (*mazG*) qPCR R | ACACCGAGACAGAAGTCAG | qRT-PCR primer for *MSMEG_5422*, reverse |
